# 3’-tRNA Fragments Target Domesticated LTR-Retrotransposons

**DOI:** 10.64898/2026.01.22.698655

**Authors:** Matthew Peacey, Joshua I. Steinberg, Andrea J. Schorn

## Abstract

Long terminal repeat (LTR) retrotransposons have been extensively co-opted by their mammalian hosts and serve essential functions. 3’-tRNA fragments (3’-tRFs) mediate post-transcriptional repression of active, murine LTR-retrotransposons through complementarity to their highly conserved tRNA primer binding site (PBS). Here, we found that 3’-tRF target sites derived from the PBS are widespread in retrotransposon-derived transcripts, suggesting that domesticated elements remain subject to regulation. Using luciferase reporters, we validated post-transcriptional repression at multiple 5’ UTR sites derived from LTR-retrotransposons. We further established paternally expressed 3 (*Peg3*), an imprinted gene with homology to retroviral Gag, as a target of an Arg-TCT 3’-tRF via a conserved 5’ UTR site. These findings provide a proof-of-principle for regulation of domesticated LTR-retrotransposons by 3’-tRFs, suggesting that their ancient role in transposon defense has been co-opted for endogenous gene regulation.

## Introduction

Transposons pose a threat to genomic integrity, but also provide the raw material for genomic innovation. Through domestication, the host can repurpose transposon-derived sequences to achieve new cellular functions, including the evolution of protein-coding genes, long non-coding RNAs (lncRNAs), and cis regulatory modules ^1,2^. Although many transposons have been co-opted in this way, long terminal repeat (LTR) retrotransposons are particularly notable for their extensive contribution to mammalian gene regulation and transcript diversity ^3^. LTR-retrotransposons mobilize via an RNA intermediate that is reverse transcribed and reintegrated in a manner analogous to retroviruses ^4^. Full-length elements consist of an internal sequence encoding retroviral proteins, flanked by 5’ and 3’ LTRs that act as promoter and termination sequences, respectively. With few exceptions, reverse transcription initiates through complementarity between the 3’ end of a specific tRNA and a primer binding site (PBS) immediately downstream of the 5’ LTR^5–7^. Elements within a family are usually constrained to the use of a specific isodecoder tRNA (i.e. a specific sequence within a group of tRNAs with identical anticodons).

LTR-retrotransposons are especially prone to co-option, in part because they contain strong promoters that can initiate transcription upstream of host genes ^8,9^. They remain regulated by transcription factors that confer tissue- or stage-specific expression patterns, particularly in early embryogenesis ^10–13^. In other cases of co-option, the internal sequence of the retrotransposon itself is incorporated into a host transcript, either as a lncRNA ^14^ or, more rarely, through direct exaptation of retroviral proteins ^15,16^. The latter is illustrated by the repeated domestication of Group-specific antigen (Gag) polyproteins, which assemble the virus-like particles of active elements but have been repurposed in mammalian proteins with diverse functions in the placenta, brain, and elsewhere ^1,16^. Because the mobile ancestors of domesticated LTR-retrotransposons were subject to host silencing, their descendants often remain influenced by that control. For example, since LTR-retrotransposons are prominent targets of DNA methylation, their domestication can give rise to imprinted loci exhibiting parent-of-origin-specific expression ^17^. Endogenous retroviruses (ERVs) closely related to infectious *Retroviridae* constitute the majority of LTR-retrotransposons in mammals, and mediate imprint establishment at lineage specific murine loci ^18,19^. By contrast, imprinted genes derived from ancient *Metaviridae* Ty3/Gypsy-elements are conserved across placental mammals ^20^.

Genome-wide epigenetic reprogramming during development transiently releases transposons from repressive chromatin ^21^. At these stages, host defense mechanisms at the RNA level become critical to restrict transposon expression and mobility ^22–24^. Small RNAs derived from the 3’ end of tRNAs (3’-tRFs) exploit complementarity to the PBS to limit the mobility of LTR-retrotransposons active in mice ^25^. 3’-tRFs are produced through endonucleolytic cleavage in the T-loop of mature tRNAs to generate fragments of 17-19 nucleotides (“tRF3a”) and 22 nucleotides (“tRF3b”) ^26,27^. Of these, tRF3a fragments interfere with reverse transcription, while tRF3b fragments post-transcriptionally silence the production of retroviral proteins ^25^. 3’-tRFs are loaded into Argonaute (AGO) proteins ^26,28–34^, and tRF3b fragments have been shown to target genes via sites in the 3’ UTR^30,32,33^. However, the full extent to which tRF3b fragments contribute to gene regulation remains poorly understood.

We propose that PBS sequences retained during LTR-retrotransposon domestication are an abundant source of tRF3b target sites in mammalian transcriptomes. To investigate such sites, we predicted 3’-tRF target sites genome-wide and analyzed their enrichment in LTR-retrotransposon derived sequences, including the 5’ UTRs of protein-coding genes. We validated several of these sites experimentally and established 3’-tRF regulation of paternally expressed 3 (*Peg3*), an imprinted gene derived from an LTR-retrotransposon. These findings provide proof-of-principle for targeting of coopted, functional retrotransposon-derived genes, expanding the known functions of 3’-tRFs beyond transposon defense ^25^ and the regulation of specific oncogenes ^30,33^.

## Results

### 3’-tRF target sites are abundant in LTR-retrotransposon derived transcripts

Building on prior evidence that 3’-tRFs repress active LTR-retrotransposons via the PBS^25,35^, we asked whether they might also recognize complementary sites within LTR-retrotransposon derived host transcripts. To identify these sites, we aligned 22 nucleotide tRF3b sequences to the mouse genome using the miRNA prediction tool miRanda ^36^ (figure 1A). Given evidence that 3’-tRFs can silence targets independently of seed pairing ^35^, we scored alignments equally across the length of the small RNA. This approach generated a genome-wide catalog of putative 3’-tRF target sites (table S1).

**Figure 1.**
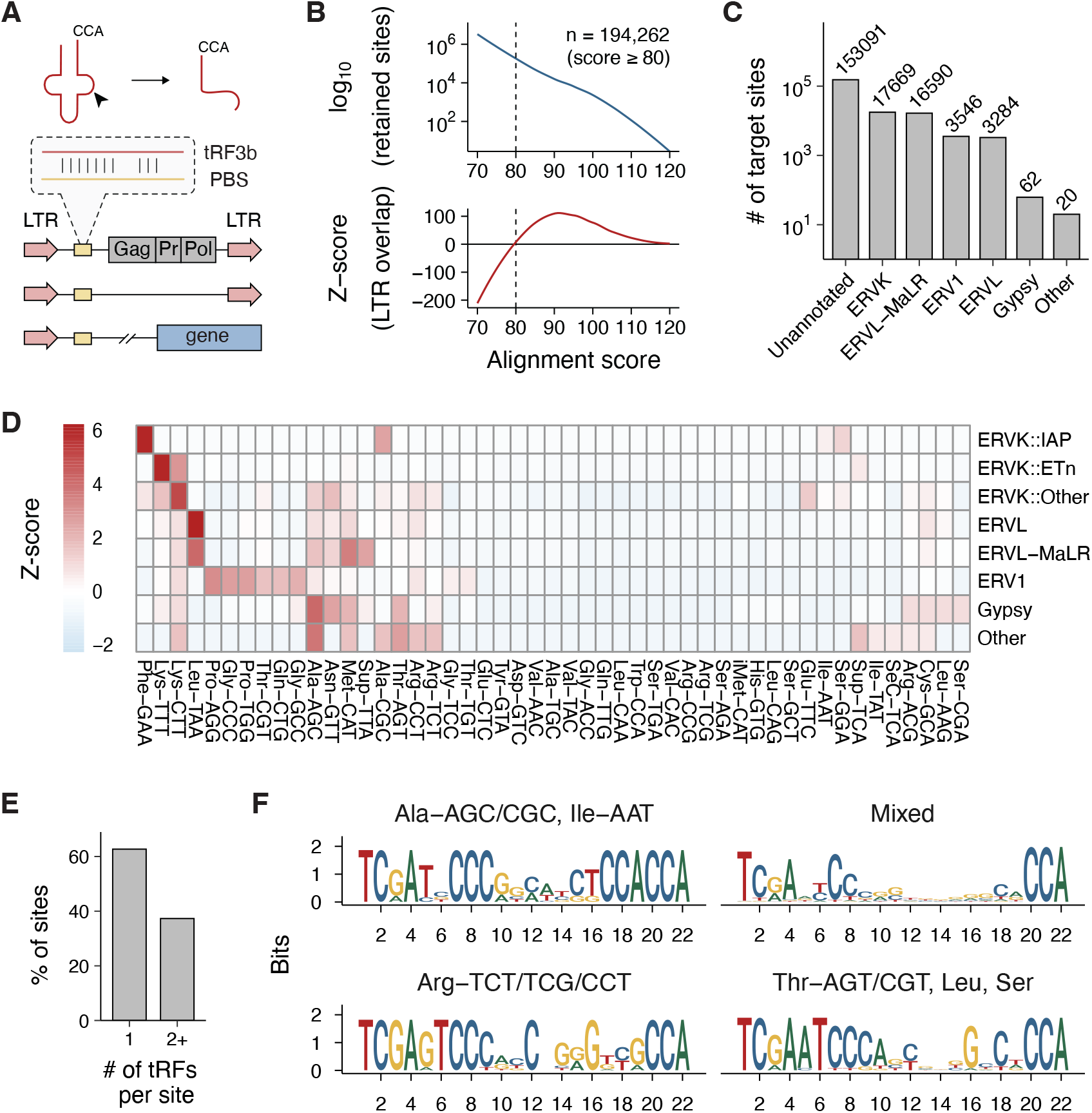
Genomic features of predicted 3’-tRF target sites. (**A**) Schematic of the target site prediction pipeline. 22 nt 3’-tRNA fragment (tRF3b) sequences were aligned to the mouse genome to identify sites corresponding to the primer binding site (PBS) of long terminal repeat (LTR) retrotransposons and to complementary sequences in host genes. (**B**) Target site number (top) and enrichment in or 200 bp downstream of LTRs (Z-score, bottom) across alignment score thresholds. A minimum score of 80 (dashed line) was used for downstream analyses. (**C**) Distribution of predicted target sites across LTR-retrotransposon families. (**D**) Heatmap showing enrichment of predicted target sites in LTR-retrotransposon sub-families (rows) by tRF3b isoacceptor (columns). Z-scores are row-scaled. (**E**) Proportion of predicted target sites with one or more than one unique tRF3b sequence aligning to the same site (alignment score *≥*70). (**F**) Sequence logos of unique tRF3b sequences grouped by sequence similarity. Hierarchical clustering of 7-mer frequency profiles using Euclidean distance resulted in four groups. The major tRNA isoacceptors contributing to each group are shown on top of the logos.

To define a meaningful alignment score cutoff, we performed a permutation analysis and selected a score threshold of 80, at which predicted target sites were significantly enriched within or downstream of LTRs relative to a randomized background (*Z* = 62, *p* = 0.01; figure 1B). Despite this enrichment, most sites above the threshold were not associated with an LTR (figure 1C). We suspect that many of these sites are LTR-retrotransposon derived, but no longer match a transposon consensus sequence due to their age and accumulated mutations. For clarity, we focused our initial analysis on target sites associated with annotated LTRs.

The distribution of 3’-tRF target sites across retrotransposon families was non-random: top scoring hits frequently matched the cognate tRNA used to prime reverse transcription of a given family (figure 1D). For example, Leu-TAA showed the strongest enrichment within the ERVL family across all 3’-tRFs. This supports the conclusion that many predicted target sites correspond to *bona fide* primer binding sites. We collapsed overlapping hits into unique genomic coordinates for downstream analysis, but noted that in 38% of cases, two or more distinct 3’-tRFs aligned to the same site (figure 1E). This likely reflects conserved sequence features at the 3’ ends of multiple tRNAs, which are apparent from clustering of 3’-tRFs by sequence similarity (figure 1F). Accordingly, in the majority of cases in which a site is hit by multiple 3’- tRFs, those 3’-tRFs originate from the same cluster (figure S1). We suspect that this may enable cooperative or redundant silencing by related 3’-tRFs.

To explore the potential regulatory impact of 3’- tRFs on host transcripts, we intersected target site coordinates with GENCODE-annotated transcripts. Most hits occurred in long non-coding RNAs, consistent with the frequent contribution of LTR-retrotransposons to these transcripts ^14^. However, target sites were underrepresented in 5’ UTRs and enriched in 3’ UTRs relative to expectation (figure 2A). We hypothesize that this pattern reflects in part cases in which a transcript initiates in an LTR with an intact PBS, but splicing to downstream exons occurs from within the LTR and hence excludes the target site from the mature transcript. To test this, we examined LTR-derived target sites within 200 bp of a 5’ splice site and found that 47.8% were excluded from the annotated 5’ UTR (figure 2B).

**Figure 2.**
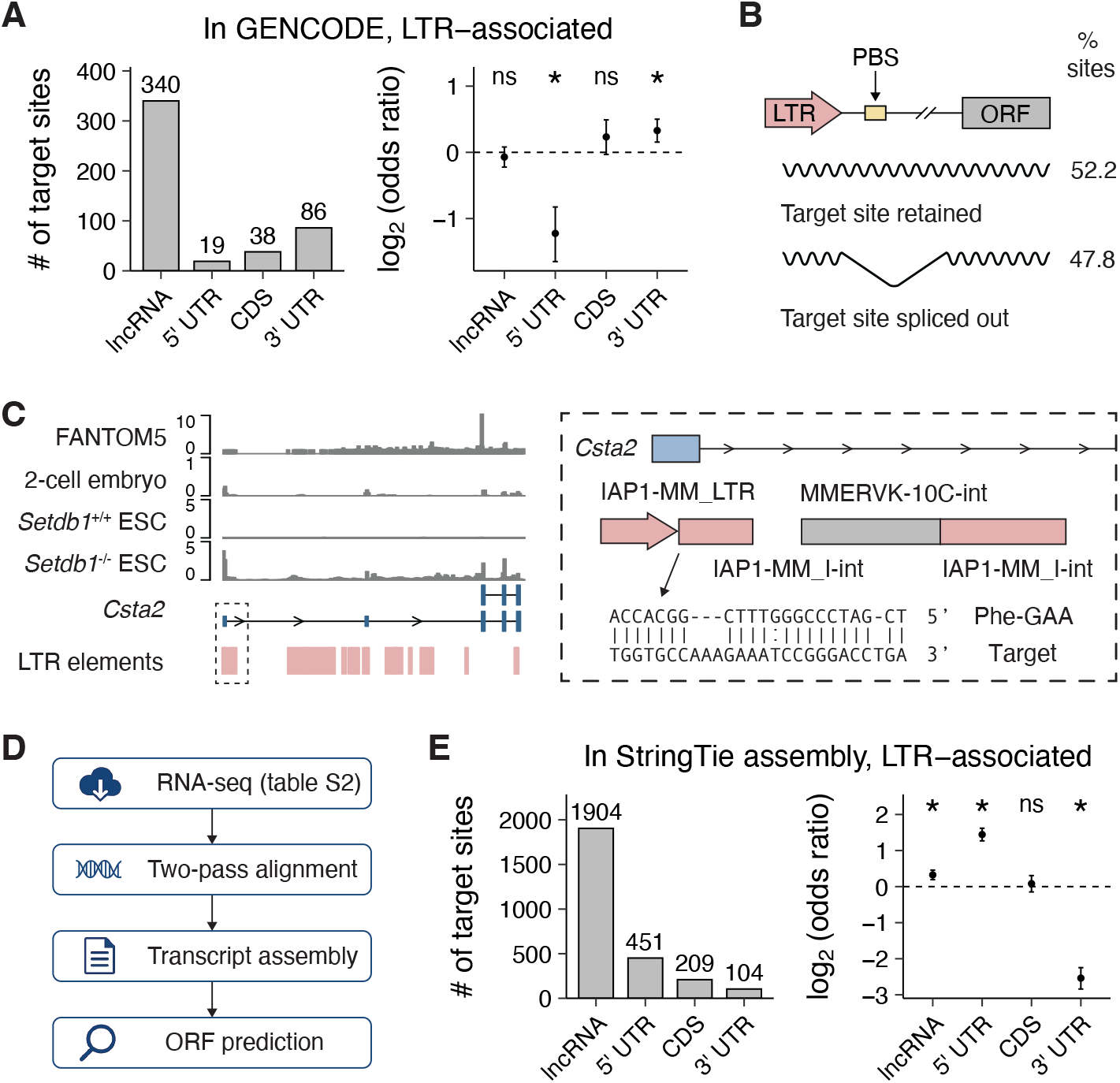
Transcriptome distribution of predicted 3’-tRF target sites. (**A**) The number of LTR-associated 3’-tRF target sites in GENCODE transcripts (left) and the enrichment at each position relative to expectation (right). Error bars show 95% confidence intervals. Asterisks (*) indicate *p*-values < 0.05 from Fisher’s exact tests. (**B**) Schematic illustrating alternative splicing outcomes at LTR-initiated protein-coding transcripts. The PBS is either retained in the 5’ UTR or spliced out, depending on the position of the 5’ splice site. Percentages reflect the proportion of LTR-associated target sites located within 200 bp upstream (retained) or downstream (spliced out) of a splice donor site. (**C**) Genome browser view of the *Csta2* locus showing an LTR-initiated transcript with a predicted 3’-tRF target site in its 5’ UTR. The FANTOM5 track shows total CAGE read counts across FANTOM5 datasets, which include diverse somatic tissues. RNA-seq tracks show expression in the 2-cell embryo (GSE66582) and in wild-type (*Setdb1*^+/+^) or *Setdb1* knockout (*Setdb1*^-/-^) mESCs (GSE29413). The start site of the LTR-retrotransposon derived transcript and the position of the predicted 3’-tRF target site are shown in detail on the right. (**D**) Schematic of the transcriptome assembly method. (**E**) The number of LTR-associated 3’-tRF target sites in assembled transcripts (left) and the enrichment at each position relative to expectation (right). Error bars show 95% confidence intervals. Asterisks (*) indicate *p*-values < 0.05 from Fisher’s exact tests.

Nonetheless, in many cases a PBS-derived 3’-tRF target site is retained in the 5’ UTR as expected. An example is the *Csta2* locus (figure 2C), which encodes a canonical transcript expressed in somatic tissues and an alternative LTR-initiated isoform expressed in the 2-cell embryo. Neither isoform is detectable in wild-type mouse embryonic stem cells (mESCs), but the isoform initiated by an IAP1 LTR is strongly expressed in *Setdb1*^-/-^ mESCs in which LTR-retrotransposons are transcriptionally de-repressed ^37,38^. Splicing from the IAP1 internal sequence retains an intact Phe-GAA primer binding site in the 5’ UTR of the resulting transcript, leading to complementarity to Phe-GAA tRF3b.

Given that 3’-tRF target sites were marginally enriched in 3’ UTRs, we tested whether such sites could support repression in a manner consistent with canonical post-transcriptional gene silencing by 3’-tRFs ^30,32–34,39^. Focusing on sites complementary to Leu-TAA tRF3b from ERVL and ERVL-MaLR elements (figure S2A), we found that a site from *Mplkipl1* (figure S2B) was sufficient to confer repression on addition of a Leu-TAA tRF3b mimic to a similar degree as a perfectly complementary site, whereas sites with additional mismatches conferred weak or no repression (figure S2C). Because repression via the 3’ UTR has been explored elsewhere ^32,33^, and because *Mplkipl1* is a strain-specific pseudogene insertion, we focused subsequent analyses on target sites in 5’ UTRs.

The GENCODE annotation primarily represents well-characterized transcripts expressed in somatic tissues, and consequently fails to capture many retrotransposon-derived transcripts with high repeat content that are expressed in specific niches such as the early embryo. Previous studies have successfully recovered such transcripts through transcriptome assembly ^12,13,38,40–44^. To investigate 3’-tRF target sites in LTR-derived transcripts absent from GENCODE, we collected RNA-seq data from the early embryo and cell culture models of early embryogenesis (table S2), assembled a transcriptome with StringTie ^45,46^, and predicted open reading frames in the assembled transcripts (figure 2D).

As for GENCODE transcripts, lncRNAs were the primary contributor to predicted 3’-tRFs targets, but in contrast to GENCODE we found an enrichment of target sites in 5’ UTRs and a depletion from 3’ UTRs (figure 2E). This likely reflects increased recovery of transcripts from intact LTR-retrotransposons with a PBS positioned upstream of a retroviral open reading frame. We also identified LTR promoter-initiated chimeric transcripts absent from GENCODE that contain a 3’-tRF target site in the 5’ UTR. One example is the *Cyp2b23* locus (figure S2D), where an RLTR9D element drives transcription that splices onto downstream exons ^38,47–49^. Importantly, splicing occurs from within the ETnERV2 internal sequence so that the transcript retains the Lys-TTT primer binding site in the 5’ UTR.

Collectively, these results indicate that 3’-tRF target sites derived from the PBS of LTR-retrotransposons are widespread in the mouse transcriptome, including in the 5’ UTRs of protein-coding genes. We next asked whether these target sites can mediate repression via their cognate 3’-tRFs.

### Functional validation of 3’-tRF target sites in the 5’ UTR

To select putative 3’-tRF target sites for experimental validation, we collected sites located within or up to 200 bp downstream of an annotated LTR that lay in the 5’ UTR of a GENCODE-annotated transcript, or the 5’ UTR of a StringTie-assembled transcript independently validated by PCR^38,48^. We also included target sites not associated with an annotated LTR, but residing in the 5’ UTR of transcripts encoding proteins homologous to retroviral Gag (table S3). For each type of target site, we adjusted the minimum alignment score threshold to restrict candidate numbers as necessary, and excluded cases in which cloning the full 5’ UTR was impractical due to length or repeat content. This resulted in a focused set of 14 candidate targets (figure 3A).

**Figure 3.**
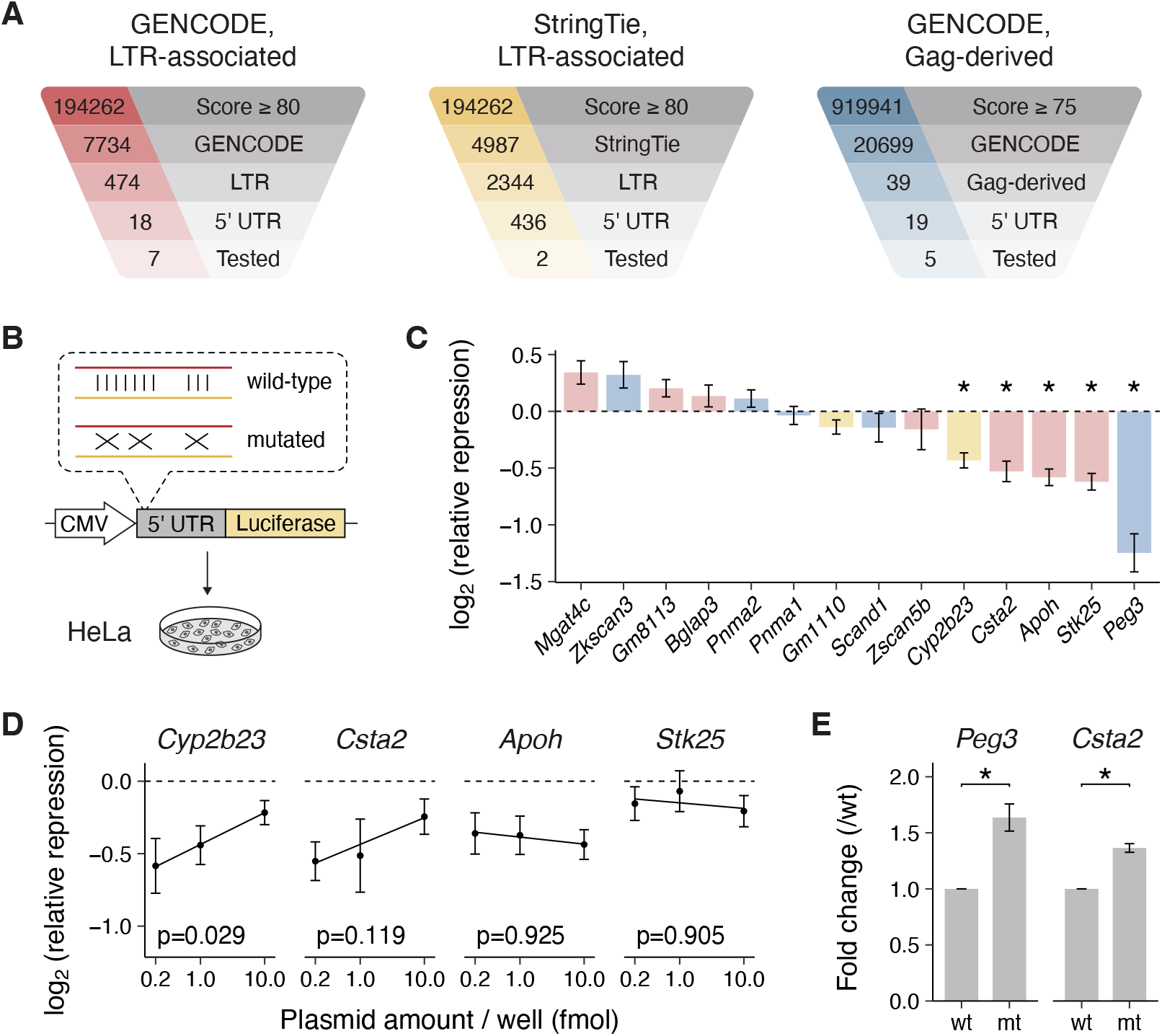
Functional validation of 3’-tRF target sites in the 5’ UTR. (**A**) Selection criteria for experimental validation of predicted 3’-tRF target sites in the 5’ UTR of protein-coding genes. Distinct filtering strategies were applied for each target site type. (**B**) Schematic of the luciferase reporter assay used to test repression via 5’ UTR target sites. (**C**) Repression of luciferase reporters containing the 5’ UTR of the indicated gene. Relative repression was calculated as the ratio of relative light units for wild-type target site reporters versus mutant. The dashed line indicates no repression. Bars are colored according to the categories used for site selection as shown in (A). Error bars show propagated standard error from technical replicates. Asterisks (*) indicate *p*-values < 0.05 from one-sided, one-sample *t* -tests. (**D**) Dose-dependent repression of luciferase reporters containing the 5’ UTR of the indicated gene. Relative repression was calculated as in (C). Error bars show propagated standard error from technical replicates. *p*-values report one-sided tests for a positive slope from a weighted linear regression. (**E**) Repression of *in vitro* transcribed luciferase reporter mRNA containing the indicated 5’ UTR. Fold change was calculated by normalizing relative light units measured for mutant (mt) and wild-type (wt) reporters. Error bars show standard error from *n* = 3 biological replicates. Asterisks (*) indicate p-values < 0.05 from one-sided, one-sample *t* -tests.

For each candidate, we cloned the full-length 5’ UTR into a luciferase reporter and compared expression of wild-type and scrambled target site variants to infer relative repression (figure 3B). HeLa cells were used because they express abundant 3’-tRFs and have previously been shown to support silencing via the 5’ UTR ^25,35^. Five candidates showed significant repression (figure 3C), which was most pronounced in the case of *Peg3*. For reporters in which repression was relatively modest, we titrated the plasmid dose and observed dose-dependence in two cases (*Cyp2b23* and *Csta2*; figure 3D), consistent with regulation by a limited pool of endogenous 3’-tRFs. Although *Cyp2b23* showed evidence of repression, the absolute luciferase signal was low relative to other reporters (figure S3A), likely due to multiple retroviral open reading frames that inhibit translation of the main ORF. Consequently, we did not pursue this candidate further. Importantly, repression of *Peg3* and *Csta2* reporters persisted when transcribed *in vitro* and transfected as mRNAs (figure 3E, S3B-C), confirming that the effect is post-transcriptional.

We chose to further explore *Peg3* as a 3’-tRF target because of the robust repression observed in luciferase assays (figure 3C, 3E), and its established role as a transcription factor regulating maternal behavior and fetal growth ^50–59^. *Peg3* is paternally imprinted and expressed primarily in the placenta, brain, and skeletal muscle ^60,61^. Although no annotated LTR is present at the *Peg3* locus, the N-terminal SCAN domain of human PEG3 is homologous to the C-terminal capsid domain of retroviral Gag, with closest similarity to the Ty3/Gypsy element *GypsyDR1* ^16,62,63^. While this homology is largely absent from mouse PEG3 as a result of loss of the SCAN domain ^64^, the 5’ UTR region containing the 3’-tRF target sites is conserved between species (figure S4A).

The 5’ UTR of *Peg3* contains two distinct regions of complementarity to 3’-tRFs: one matching Ala-AGC 3’-tRF and one matching several 3’-tRFs (sites “A” and “B”, respectively; figure 4A). In the initial reporter assays, both sites were mutated simultaneously (figure 4B, variant 1). To determine their respective contributions, we mutated each site individually and found that site A had no effect on luciferase output (figure 4B, variant 2), whereas mutation of site B alone fully recapitulated derepression of the double mutant (figure 4B, variants 3 and 4). Moreover, repression at site B is specific to the 3’-tRF complementary region, because mutation of the immediately adjacent bases had no effect (figure 4B, variant 5).

**Figure 4.**
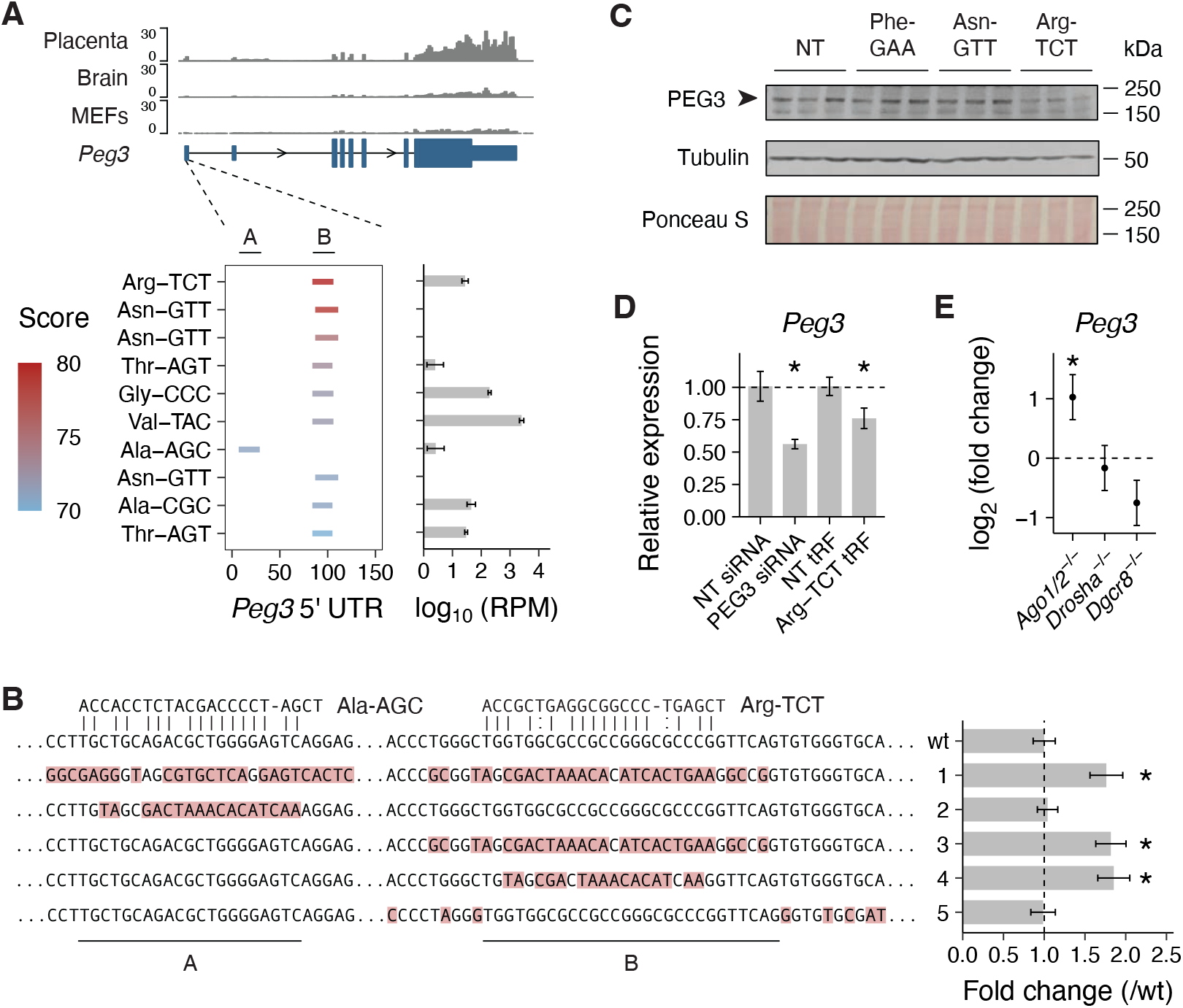
*Peg3* is a target of Arg-TCT 3’-tRF. (**A**) Genome browser view of the murine *Peg3* locus. RNA-seq tracks show expression in placenta (ENCFF516KLL), E14.5 whole brain (ENCFF570RBK), and embryonic fibroblasts (ENCFF918FWL). The *Peg3* 5’ UTR is enlarged with predicted 3’-tRF target sites colored by alignment score. For each 3’-tRF, abundance in HeLa cells estimated from small RNA-seq data (GSE82199) is shown in reads per million (RPM). Error bars show standard error from *n* = 3 biological replicates. (**B**) Luciferase reporter assay in HeLa cells testing *Peg3* 5’ UTR target site variants. Fold change was calculated by normalizing relative light units measured for each target site variant to those of the wild-type (wt) reporter. Red highlighting indicates mutated nucleotides in each variant. The alignment of top scoring tRF3b sequences at each site is shown above the wild-type sequence. Error bars show propagated standard error from technical replicates. Asterisks (*) indicate *p*-values < 0.05 from one-sided, one-sample *t* -tests. (**C**) Western blot showing endogenous PEG3 expression in mouse embryonic fibroblasts transfected with the indicated tRF3b mimics. Probing for tubulin and Ponceau S staining served as loading controls. (**D**) *Peg3* mRNA levels in P19 embryonal teratocarcinoma cells transfected with the indicated siRNAs or tRF3b mimics. Expression was calculated relative to samples transfected with a non-targeting (NT) siRNA. Error bars show propagated standard error from technical replicates. Asterisks (*) indicate *p*-values < 0.05 from one-sided, one-sample *t* -tests. (**E**) *Peg3* mRNA levels in mESCs of the indicated genotype. Fold change was calculated relative to wild-type. Error bars show standard error. Asterisks (*) indicate *p*-values < 0.05 from differential expression analysis. Data are from GSE122627, GSE110942, GSE78971 and GSE78974.

These findings suggest that site B alone contributes to repression of *Peg3*, which is consistent with its conservation across species, whereas site A is mouse specific (figure S4A). Of the top scoring 3’-tRFs at site B, Arg-TCT tRF3b is moderately expressed in HeLa cells and contiguously base pairs to the central region of the target site. By contrast, Asn-GTT tRF3b is sparsely expressed and pairs to the target site through nucleotides 2-6, consistent with miRNA-like seed pairing (figure S4B). None of the 3’-tRFs aligning to site B matched priming tRNAs that we could assign to the Ty3/Gypsy elements *GypsyDR1* and *MDG-1* (figure S4C). Similarly, various 3’-tRFs were the top scoring hits at annotated mouse Gypsy elements (figure 1D), but very few corresponded to those predicted to bind site B. Whether site B is derived from a PBS therefore remains unclear, although the diversity of tRNA primers among Gypsy-family elements ^5,65^ makes such an origin plausible.

To confirm a direct role for 3’-tRFs in silencing of endogenous *Peg3*, we transfected mouse embryonic fibroblasts with synthetic 3’-tRF mimics and observed downregulation of PEG3 protein specific to Arg-TCT tRF3b (figure 4C). The same mimic mildly reduced *Peg3* RNA levels in P19 embryonal teratocarcinoma cells (figure 4D). Finally, re-analysis of RNA-seq data ^66^ showed that *Peg3* RNA is elevated in *Ago1/2*^-/-^ mESCs, but not in *Drosha*^-/-^ or *Dgcr8*^-/-^ cells lacking most miRNAs (figure 4E), consistent with post-transcriptional small RNA-mediated repression independent of miRNAs. Together, these data suggest that *Peg3* is a target of Arg-TCT 3’-tRF via a 5’ UTR site of LTR-retrotransposon origin. Such target sites are widely distributed in mammalian transcriptomes and may represent a broader mechanism of post-transcriptional regulation affecting genes with developmental functions.

## Discussion

We identified thousands of potential target sites for 3’- tRFs based on sequence complementarity using an approach that allowed for gaps, which are common in *bona fide* primer binding sites ^67^, but that did not enforce seed requirements, which do not necessarily apply to 3’-tRFs ^35^. By intersecting these target sites with a transcriptome annotation from GENCODE or assembled from RNA-seq data, we uncovered abundant examples in LTR-retrotransposon derived transcripts. Many of these resemble primer binding sites in their position within annotated LTR-retrotransposons and the extent of complementarity to cognate 3’-tRFs. Using a luciferase reporter approach, we validated repression of candidates derived from LTR-retrotransposons with 3’-tRF target sites in their 5’ UTR. We further established *Peg3* as a proof-of-principle for a novel mode of post-transcriptional regulation affecting developmentally expressed genes, which may have widespread consequences in health and disease.

Our prediction strategy is conceptually similar to that used by tRFtarget ^68,69^, although we note that their collection of 3’-tRF sequences is incomplete. Specifically, the primary source of mouse 3’-tRFs is tRFdb ^70^, which explicitly filters out multi-mapping tRFs and therefore excludes many that target repeat sequences. As a result, prominent targets derived from LTR-retrotransposons are missed by tRFtarget. Unlike other prediction efforts that utilize functional data from somatic cell lines ^71–74^, we aligned 3’-tRF sequences genome-wide with the aim of capturing all LTR-retrotransposon derived targets. This approach captures contexts in which LTR-retrotransposons are transiently de-repressed, in particular early embryogenesis, and includes target sites in the 5’ UTR, consistent with the expected position of the PBS. We considered incorporating *in vivo* AGO2 CLIP data ^75^ to refine our prediction, but found that 3’-tRFs are sparsely represented in these libraries. In addition, AGO2 binding does not necessarily correlate with miRNA-3’ UTR repression ^76^. Transcriptomic validation strategies successfully applied to miRNAs ^66,76,77^ are limited for 3’-tRFs, owing to our incomplete knowledge of specific biogenesis or effector factors beyond the recently identified 2’O-methyltransferase HENMT1^35^. As CLIP datasets expand to include additional AGO family members and further 3’-tRF specific factors are characterized, these approaches should enable high-throughput target discovery in physiological contexts.

Targeting rules for repression by 3’-tRFs are emerging, but have been limited to specific sequence contexts. Prior work focused on target sites in the 3’ UTR has suggested that tRF3b follow miRNAs in their dependence on a seed sequence at nucleotides 2-6 of the small RNA^30,33^. However, a recent massively parallel reporter assay from our lab examining a Lys-TTT 3’-tRF target site in the 5’ UTR of the MusD retrotransposon showed that repression is seed-independent ^35^. Consistent with this, Arg-TCT tRF3b has a poor seed match to the *Peg3* 5’ UTR but contiguously base pairs at its center, reminiscent of the “centered” sites reported for specific miRNAs ^78^. Conversely, Asn-GTT tRF3b has a 6-mer seed match supplemented by extensive 3’ terminal complementarity (figure S4B), yet fails to repress *Peg3* (figure 4C). Similarly, the Leu-TAA 3’-tRF target site in *Mplkipl1* has a mismatch at position 2 and a two-nucleotide bulge in the seed region (figure S2B), yet supports roughly two-fold repression by a Leu-TAA tRF3b mimic (figure S2C). Notably, primer binding sites for active LTR-retrotransposons are typically 18 nucleotides or shorter ^5^, which may explain why 3’-tRFs have evolved to repress these elements and their domesticated ancestors without dependence on the seed.

Based on the position of the PBS immediately downstream of the LTR promoter, 3’-tRF target sites in LTR-retrotransposon derived genes should occur predominantly in the 5’ UTR. The relative depletion of 5’ UTR sites in GENCODE transcripts likely reflects a combination of PBS decay through genetic drift, transcription from solo LTRs that lack downstream internal transposon sequence, and the use of splice donor sites upstream of the PBS that exclude it from the mature transcript (figure 2B). PBS-like target sites in the 3’ UTR could arise from tRNA-derived sequences, including tRNA-derived SINE elements that are overrepresented in the 3’ UTR of mouse transcripts ^9^. Alternatively, an LTR-retrotransposon can lie immediately downstream of an open reading frame, as is the case for *Mplkipl1* in the reference genome strain (figure S2B). In these cases, incorporation of transposon sequence into the 3’ UTR would require read-through transcription across the entire LTR sequence despite its inherent poly(A)-signal. The fact that a Leu-TAA 3’-tRF target site derived from the MERVL PBS can support repression of a reporter (figure S2C) suggests that more MERVL-derived target sites may be functional and warrant further investigation.

The intersection of predicted 3’-tRF target sites with GENCODE and RepeatMasker annotations identifies many LTR-retrotransposon derived targets (figure 2A), but likely misses those at the extremes of evolutionary age. On the one hand, young, intact elements are well represented in RepeatMasker but often excluded from GENCODE. Transcriptome assembly from early embryo RNA-seq data recovers some examples (figure 2E), but further investigation is difficult without validation of transcript structure. Further confidence can be gained by defining genuine transcription start sites, which has recently been addressed in the early mouse embryo via Smart-seq+5’ ^40^, although reliance on oligo-dT or random priming during standard cDNA generation often leads to low coverage of 5’ RNA ends. Techniques that enrich for 5’-ends of capped mRNAs have revealed the full extent of transposon-initiated transcript isoforms ^9,79^, and are being developed to capture low input, embryonic samples ^80^. Ultimately, confidence in individual target transcripts requires locus-by-locus validation of transcript structure and function. Such validation is warranted by numerous reports of functional LTR-retrotransposon derived isoforms specific to the early embryo ^11–13,81^.

At the opposite extreme, deeply domesticated loci are poorly annotated in RepeatMasker yet have the greatest potential for functional integration into host genes. This is exemplified by *Peg3*, the retrotransposon origin of which was first recognized through homology of the SCAN domain to Ty3/Gypsy Gag in human ^16,62,63^. The mouse orthologue has since lost overt homology to Gag, but retains a conserved and functional 3’-tRF target site in its 5’ UTR, likely derived from the PBS of the Ty3/Gypsy element. We expect that many additional 3’-tRF target sites of latent origin in LTR-retrotransposons remain to be identified.

The effects of 3’-tRFs on gene output are comparable in magnitude to miRNA-mediated repression and thus suited to the fine-tuning of dosage-sensitive genes ^82,83^. These may include loci transiently expressed during developmental transitions, which are frequently co-opted from LTR-retrotransposons in early embryogenesis ^84^. Imprinted genes such as *Peg3* represent a special case of dosage sensitivity, since their monoallelic expression implies a selective constraint on transcript abundance ^85^. Speculatively, allele-specific expression of 3’-tRF biogenesis factors could enable 3’-tRFs to reinforce these imprints, analogous to the reciprocal regulation of the paternally imprinted gene retrotransposon Gaglike 1 (*Rtl1*) by maternally expressed miRNAs ^86,87^. In addition to regulating genes that support the normal progression of development, 3’-tRFs may repress aberrant LTR-initiated transcripts that would otherwise produce truncated or mis-expressed proteins, as occurs in tumorigenesis ^88^. Conversely, cancer cells may exploit LTR promoters that lack a primer binding site to allow an oncogene to evade 3’-tRF mediated repression.

LTR-retrotransposons active in mice are restricted by 3’- tRFs through complementarity to their primer binding site ^25^. The discovery of analogous 3’-tRF target sites in LTR-retrotransposon derived host transcripts suggests that this ancient defense mechanism has been co-opted for endogenous gene regulation. The repurposing of transposon defense mechanisms is a recurrent theme in mammalian evolution, exemplified by the domestication of KRAB zinc-finger proteins for transcriptional control ^89^, SPEN targeting of *Xist* through an LTR-derived repeat ^90^, and piRNA-mediated silencing of an LTR-initiated transcript at the *Rasgrf1* locus ^91^. Our findings suggest a broader impact of 3’-tRF mediated regulation on gene expression programs in development and disease, shedding light on how ancient retroviral sequences continue to shape host genome function.

## Materials and methods

### Cell culture

All cell lines were maintained at 37°C in 5% CO_2_ and sub-cultured using 0.25% trypsin-EDTA (Gibco; 25200056) diluted 1:1 with PBS pH 7.2 (Gibco; 20012027). HeLa (ATCC; CCL-2), HEK293T (ATCC; CRL-3216), and mouse embryonic fibroblasts (ATCC; SCRC-1008) were grown in DMEM (Corning; 10-013-CV) with 10% fetal bovine serum (Corning; 35-010-CV). P19 embryonal teratocarcinoma cells (ATCC; CRL-1825) were grown in MEM α (Gibco; 12571063) with 2.5% fetal bovine serum and 7.5% bovine calf serum (VWR; 10158-358) or, for P19 RNA-seq, 10% fetal bovine serum only.

### Prediction of 3’-tRF target sites

Scripts used for target prediction are available at the project [GitHub repository](https://github.com/mwpeacey/tRF_target_prediction). High confidence *Mus musculus* (mm10) mature tRNA sequences were downloaded from [GtRNAdb](https://gtrnadb.org/genomes/eukaryota/Mmusc10/mm10-mature-tRNAs.fa) ^92^ (release 22), “CCA” appended to each, and the terminal 22nt extracted to define the 3’-tRF sequence. Unique sequences were assigned numeric identifiers and written to a FASTA file. The mm10 primary genome assembly was downloaded from [GENCODE](https://ftp.ebi.ac.uk/pub/databases/gencode/Gencode_mouse/release_M23/GRCm38.primary_assembly.genome.fa.gz) and split into 10 kbp windows, overlapping by 50 bp, to accommodate memory restrictions during alignment. For each 3’-tRF sequence, genomic target sites were identified using miRanda ^36^ (v1.9) with seed-weighting removed (miranda -sc 70.0 -en 0.0 -scale 1.0 -loose). Alignment scores and genomic coordinates were aggregated and written to .bed and .csv formats.

### Features of genomic target sites

Predicted target sites were annotated using a custom R script and exported to .csv format (table S1). To remove redundancy from overlapping predictions, target sites were grouped by genomic interval, and only the highest-scoring tRF was retained per group. The RepeatMasker annotation (updated April 8th 2021) was downloaded from the [UCSC Table Browser](https://genome.ucsc.edu/cgi-bin/hgTables), filtered for LTR-class repeats, and internal portions (“-int”) excluded. Overlaps with target sites were defined by extending each element 200 bp downstream, since PBS sequences are typically a few nucleotides downstream of the LTR.

Enrichment of predicted target sites within or downstream of LTRs was assessed using permutation testing via regioneR ^93^. At alignment score cutoffs in 5-point increments, the observed intersection of non-overlapping target sites with LTRs (or 200bp downstream) was compared to a null distribution generated from 100 randomizations across the mm10 genome.

To generate a heatmap of the top scoring 3’-tRFs at LTR-associated target sites, “ERVK” family elements were further divided into “IAP”, “ETn” and “other” sub-families, based on the identity of the internal sequence associated with each LTR. In cases where a 3’-tRF could be derived from more than one tRNA isoacceptor, a single anticodon was selected arbitrarily.

To identify sequence patterns among 3’-tRFs, 7-mer frequencies were computed using Biostrings and clustered into four groups by hierarchical clustering (Euclidean distance, Ward’s method). Sequence logos sites of latent origin in LTR-retrotransposons remain to be identified.

The effects of 3’-tRFs on gene output are comparable in magnitude to miRNA-mediated repression and thus suited to the fine-tuning of dosage-sensitive genes ^82,83^. These may include loci transiently expressed during developmental transitions, which are frequently co-opted from LTR-retrotransposons in early embryogenesis ^84^. Imprinted genes such as *Peg3* represent a special case of dosage sensitivity, since their monoallelic expression implies a selective constraint on transcript abundance ^85^. Speculatively, allele-specific expression of 3’-tRF biogenesis factors could enable 3’-tRFs to reinforce these imprints, analogous to the reciprocal regulation of the paternally imprinted gene retrotransposon Gaglike 1 (*Rtl1*) by maternally expressed miRNAs ^86,87^. In addition to regulating genes that support the normal progression of development, 3’-tRFs may repress aberrant LTR-initiated transcripts that would otherwise produce truncated or mis-expressed proteins, as occurs in tumorigenesis ^88^. Conversely, cancer cells may exploit LTR promoters that lack a primer binding site to allow an oncogene to evade 3’-tRF mediated repression.

LTR-retrotransposons active in mice are restricted by 3’- tRFs through complementarity to their primer binding site ^25^. The discovery of analogous 3’-tRF target sites in LTR-retrotransposon derived host transcripts suggests that this ancient defense mechanism has been co-opted for endogenous gene regulation. The repurposing of transposon defense mechanisms is a recurrent theme in mammalian evolution, exemplified by the domestication of KRAB zinc-finger proteins for transcriptional control ^89^, SPEN targeting of *Xist* through an LTR-derived repeat ^90^, and piRNA-mediated silencing of an LTR-initiated transcript at the *Rasgrf1* locus ^91^. Our findings suggest a broader impact of 3’-tRF mediated regulation on gene expression programs in development and disease, shedding light on how ancient retroviral sequences continue to shape host genome function.

## Materials and methods

### Cell culture

All cell lines were maintained at 37°C in 5% CO_2_ and sub-cultured using 0.25% trypsin-EDTA (Gibco; 25200056) diluted 1:1 with PBS pH 7.2 (Gibco; 20012027). HeLa (ATCC; CCL-2), HEK293T (ATCC; CRL-3216), and mouse embryonic fibroblasts (ATCC; SCRC-1008) were grown in DMEM (Corning; 10-013-CV) with 10% fetal bovine serum (Corning; 35-010-CV). P19 embryonal teratocarcinoma cells (ATCC; CRL-1825) were grown in MEM α (Gibco; 12571063) with 2.5% fetal bovine serum and 7.5% bovine calf serum (VWR; 10158-358) or, for P19 RNA-seq, 10% fetal bovine serum only.

### Prediction of 3’-tRF target sites

Scripts used for target prediction are available at the project GitHub repository. High confidence *Mus musculus* (mm10) mature tRNA sequences were downloaded from GtRNAdb ^92^ (release 22), “CCA” appended to each, and the terminal 22nt extracted to define the 3’-tRF sequence. Unique sequences were assigned numeric identifiers and written to a FASTA file. The mm10 primary genome assembly was downloaded from GENCODE and split into 10 kbp windows, overlapping by 50 bp, to accommodate memory restrictions during alignment. For each 3’-tRF sequence, genomic target sites were identified using miRanda ^36^ (v1.9) with seed-weighting removed (miranda -sc 70.0 -en 0.0 -scale 1.0 -loose). Alignment scores and genomic coordinates were aggregated and written to .bed and .csv formats.

### Features of genomic target sites

Predicted target sites were annotated using a custom R script and exported to .csv format (table S1). To remove redundancy from overlapping predictions, target sites were grouped by genomic interval, and only the highest-scoring tRF was retained per group. The RepeatMasker annotation (updated April 8th 2021) was downloaded from the UCSC Table Browser, filtered for LTR-class repeats, and internal portions (“-int”) excluded. Overlaps with target sites were defined by extending each element 200 bp downstream, since PBS sequences are typically a few nucleotides downstream of the LTR.

Enrichment of predicted target sites within or downstream of LTRs was assessed using permutation testing via regioneR ^93^. At alignment score cutoffs in 5-point increments, the observed intersection of non-overlapping target sites with LTRs (or 200bp downstream) was compared to a null distribution generated from 100 randomizations across the mm10 genome.

To generate a heatmap of the top scoring 3’-tRFs at LTR-associated target sites, “ERVK” family elements were further divided into “IAP”, “ETn” and “other” sub-families, based on the identity of the internal sequence associated with each LTR. In cases where a 3’-tRF could be derived from more than one tRNA isoacceptor, a single anticodon was selected arbitrarily.

To identify sequence patterns among 3’-tRFs, 7-mer frequencies were computed using Biostrings and clustered into four groups by hierarchical clustering (Euclidean distance, Ward’s method). Sequence logos were generated for each of the resulting clusters using ggseqlogo.

### P19 RNA-seq

P19 cells (150,000 per well) were seeded in 12-well plates one day prior to transfection with 1 μg pmaxGFP (Lonza) using Lipofectamine 2000 (Invitrogen; 11668027). One day after transfection, cells were expanded to 6-well plates and on day three harvested in TRIzol (Thermo Fisher Scientific; 15596026). 500 pg total RNA was used as input for the SMARTer Stranded Total RNA-Seq Pico Input Kit v2 (Takara; 634412) or the NEBNext Single Cell/Low Input RNA Library Prep Kit (New England Biolabs; E6420L) to generate random-primed and poly(A)-primed RNA-seq libraries, respectively. Libraries were quantified by Qubit (Thermo Fisher Scientific; Q32851) and sequenced in a NextSeq500 (Illumina) PE76 run. Data was used for transcriptome assembly as described below.

### Transcriptome assembly and annotation

The GENCODE vM23 annotation was downloaded via AnnotationHub and target sites overlapping transcript features were identified using GenomicFeatures ^94^, assigned in priority of 5’ UTR, 3’ UTR, CDS, and lncRNA exons. Because the target prediction was performed on the genome, this approach will exclude target sites overlapping splice junctions. For LTR-initiated protein-coding transcripts, the distance from each target site to the nearest 5’ splice site was calculated to assess whether splicing excluded the site from the annotated 5’ UTR.

For transcriptome assembly, RNA-seq data listed in table S2 were downloaded from the sequence read archive using sra-tools (v3.0.9). Adapters, poly(A) tails, and low-quality bases were removed using cutadapt ^95^ (v4.5; cutadapt -a AGATCGGAA-GAGCACACGTCTGAACTCCAGTCA -A AGATCGGAA-GAGCGTCGTGTAGGGAAAGAGTGT –poly-a –quality-cutoff=15,10 –minimum-length=25). For Takara P19 libraries, 3 bases were additionally removed from the 5’ end of read 2 (cutadapt -U 3). The *Mus musculus* (mm10) primary genome assembly and the vM23 transcriptome annotation were downloaded from GENCODE and an index generated using STAR^96^ (v2.7.11a; STAR – runMode genomeGenerate –sjdbOverhang 100). Reads were aligned in two-passes as described by Modzelewski *et al*. 2021^12^ (STAR –outFilterMultimapNmax 100 – winAnchorMultimapNmax 200 –chimMultimapNmax 100). Transcripts were assembled using StringTie ^45,46^ (v3.0.0; stringtie -c 2 -f 0.05 -j 2 -s 5) and merged into a unified GTF. FASTA sequences were extracted from the merged transcriptome, and open reading frames predicted using orfipy ^97^ (v0.0.4; orfipy –strand f –start ATG –min 300). Predicted ORFs were matched to the mouse Refseq database using blast ^98^ (v2.16.0; blastp -max_target_seqs 1). For transcripts with multiple predicted ORFs, only that with the highest bitscore was retained. Target sites were assigned to these transcripts as described for GENCODE transcripts.

Genes with protein homology to retroviral Gag (table S3) were collected from Campillos *et al*. 2006^16^, supplemented with a search for “Gag” in the [Mouse Genome Informatics](https://www.informatics.jax.org/quicksearch/summary?queryType=exactPhrase&query=Gag&submit=Quick%0D%0ASearch) database.

### Cloning

PCR-amplified fragments were generated with Q5 High-Fidelity DNA Polymerase (NEB; M0491) using primers as indicated in table S4. Reaction products were purified by column cleanup (Qiagen; 28104) or gel extraction (Qiagen; 28704). Ligation-independent cloning was performed with NEBuilder HiFi DNA Assembly Master Mix (NEB; E2621), and ligations with T4 DNA ligase (NEB; M0202). Plasmids were propagated in NEB Stable Competent *E*.*coli* (NEB; C3040) and prepared using ZymoPURE plasmid kits (Zymo Research; D4208T, D4200, and D4202). All sequences that underwent PCR amplification or DNA synthesis were verified by sequencing.

### 5’ UTR luciferase reporters

The pCMV-firefly luciferase vector was generated by PCR amplification of the CMV promoter from pCMV-MusD6^99^ and insertion into pGL4.21 (Promega; E6761) via XhoI/HindIII digestion. The 5’ UTR of each candidate target, containing either an intact or scrambled target site, was synthesized as a gene fragment (Integrated DNA Technologies, table S5) and cloned into HindIII/PspOMI-digested pCMV-firefly by ligation independent-cloning. Variants of the *Peg3* pCMV-firefly reporter were generated by insertion of a PCR-amplified oligo pool (Integrated DNA Technologies, table S4) into a PCR-amplified backbone by ligation-independent cloning. HeLa cells (75,000 per well) were seeded in 24-well plates one day prior to transfection with 1 fmol (*≈*4 ng) firefly reporter, or otherwise the amount indicated, and 10 fmol (26 ng) renilla control plasmid pGL4.74 (Promega; E6921) using Lipofectamine 2000 (Invitrogen; 11668027).

*In vitro* transcribed reporters were generated by PCR amplification of the 5’ UTR from *Peg3* and *Csta2* pCMV-firefly reporters, and insertion into NheI-digested psiCheck2-XNS (Addgene; 196655) by ligation-independent cloning. Reporter RNA was synthesized using the mMESSAGE mMACHINE T7 Transcription Kit (Invitrogen; AM1344) and MEGAclear Transcription Clean-Up Kit (Invitrogen; AM1908) on XhoI-digested template. Product length and integrity were checked using an RNA ScreenTape (Agilent; 5067-5576) and TapeStation (Agilent). HeLa cells (150,000 per well) were seeded in 12-well plates one day prior to transfection with 100 ng reporter RNA using Lipofectamine MessengerMAX (Invitrogen; LMRNA008).

### 3’ UTR luciferase reporters

Annealed and phosphorylated oligonucleotides (Integrated DNA Technologies; table S4) were ligated into XhoI/NotI digested psiCheck2-XNS (Addgene; 196655). HEK293T cells (100,000 per well) were seeded 24-well plates one day prior to transfection with 2.5 fmol (*≈*10 ng) psiCheck2 reporter and 50 pmol synthetic 3’-tRF mimics (Integrated DNA technologies, 5’ phosphorylated, table S4) using Lipofectamine 2000 (Invitrogen; 11668027).

### Luciferase assays

Luciferase assays were performed with the DualGlo Luciferase Assay System (Promega; E2940) 24 hours post-transfection. Luminescence was measured using a GloMax Discover microplate reader (Promega; GM3000) with a 0.3 second integration time. For DNA reporters, cells were lysed in a 1:1 mixture of Dual-Glo reagent and PBS pH 7.2 (Gibco; 20012027), and relative light units (RLUs) were calculated as the ratio of test and internal control luminescence readings. For RNA reporters, cells were lysed with Glo lysis buffer (Promega; E2661) and RLUs were calculated by normalizing luminescence readings to total protein concentration, measured by Pierce BCA protein assay (Thermo Scientific; 23225). Relative repression by endogenous 3’-tRFs was calculated by normalizing the RLU value for wild-type constructs to that of the mutant. Relative repression by synthetic 3’-tRF mimics was calculated in two steps: first, the RLU of each reporter was normalized to that of the empty psiCheck2-XNS reporter; second, these values were expressed relative to the matched non-targeting control 3’-tRF condition. Thus, for reporter *r* and mimic *m*:

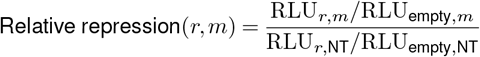

### PEG3 silencing by 3’-tRF mimics

Mouse embryonic fibroblasts (200,000 cells per well) were seeded in 6-well plates one day prior to transfection with 187 pmol synthetic 3’-tRF mimics (Integrated DNA technologies, 5’ phosphorylated, table S4) using Lipofectamine RNAiMAX (Invitrogen; 13778075). Protein was harvested 48 hours post-transfection. P19 cells (150,000 per well) were seeded in 12-well plates and the next day transfected with 30 pmol ON-TARGETplus siRNAs (Horizon Discovery, D-001810-10-05 and L-040038-01-0005) or 90 pmol synthetic 3’-tRF mimics (Integrated DNA technologies, 2’O methylated at every position, table S4). RNA was harvested 48 hours post-transfection.

### Western blots

Cells were washed with ice-cold PBS pH 7.2 (Gibco; 20012027) and lysed in ice-cold RIPA buffer (150 mM NaCl, 1% NP-40, 0.1% SDS, 0.5% sodium deoxycholate, 50 mM Tris-Cl pH 8.0) with 1X cOmplete mini EDTA-free protease inhibitor (Roche; 11836170001). Cells were scraped from the dish surface and sonicated for 5 minutes in 30 second on/off cycles. Cell debris was pelleted by centrifuging at 18,000 rcf for 10 minutes at 4°C. Proteins were precipitated from the supernatant in 80% acetone, and resuspended in a minimum volume of 1X reducing agent-free Laemmli buffer (50 mM Tris-Cl pH 6.8, 10% glycerol, 1% SDS). Total protein was quantified using the Pierce BCA protein assay (Thermo Scientific; 23225).

DTT (50 mM) and bromophenol blue (0.025% w/v) were added to the lysates before they were heated for 5 minutes at 95°C and centrifuged at 18,000 rcf for 5 minutes at 4°C. Proteins were separated by 7.5% SDS-PAGE and transferred onto a 0.45 μM nitrocellulose membrane (Bio-Rad; 1620115) using the Mini Trans-Blot system (Bio-Rad; 1703930). Transfer efficiency was checked by staining with Ponceau S (0.1% w/v) in acetic acid (5% v/v). Membranes were blocked in 5% Blotto non-fat dry milk (Santa Cruz Biotechnology; sc-2324) in TBS with 0.1% Tween 20 (Thermo Scientific; J20605-AP) for a minimum of 1 hour. All antibodies were diluted in the same buffer, used as follows: anti-PEG3 (Thermo Scientific; PA5-99683) 1:500 overnight at 4°C; anti-tubulin (Abcam; ab6046) 1:1000 for 1 hour at room temperature; anti-rabbit-HRP (Jackson Immuno Research; 111-035-144) 1:3000 for 1 hour at room temperature. Blots were developed using Pierce ECL Western Blotting Substrate (Thermo Scientific; 32106) and an Odyssey FC Imaging System (LICORbio).

### RNA extraction and RT-qPCR

RNA was extracted using TRIzol (Thermo Fisher Scientific; 15596026) with 80% v/v ethanol washes during precipitation. Total RNA was treated with TURBO DNaseI (Thermo Fisher Scientific; AM1907). Reactions were stopped by heat inactivation after addition of EDTA and cleaned up using RNAClean XP beads (Beckman Coulter; A63987). RNA was reverse transcribed using Superscript III (Thermo Fisher Scientific; 18080044) and quantified using PowerTrack SYBR Green Master Mix (Thermo Fisher Scientific, A46109). Reactions were run on a QuantStudio 6 Flex Real-Time PCR System (Applied Biosystems; 4485691). Relative abundance was calculated relative to non-targeting 3’-tRF using the ΔΔCt method, using Gapdh as an internal control. All primers used are listed in table S4.

### *Peg3* alignment

The sequence of the first exon of the canonical *Peg3* transcript in each species was downloaded from Ensembl and aligned in R using the ClustalOmega method, via msa ^100^ (v1.40.0). Alignments were plotted using ggmsa ^101^ (v1.14.1).

### Primer tRNA analysis

To predict priming tRNAs of elements ancestral to *Peg3*, the terminal 18nt of mm10 mature tRNAs were extracted as described for 22nt sequences in “Prediction of 3’-tRF target sites”. The *Danio rerio* (danRer11) and *Drosophila melanogaster* ([dm6](https://hgdownload.soe.ucsc.edu/goldenPath/dm6/bigZips/)) primary genome assemblies were downloaded from UCSC and an index built with bowtie ^102^ (v1.2.1.1). 18nt 3’-tRF sequences were aligned to each genome, allowing up to 3 mismatches and reporting all alignments (bowtie -f -a -v 3). Aligned sequences were intersected with the respective RepeatMasker annotation downloaded from the UCSC Table Browser.

### Re-analysis of published data

RNA-seq data displayed on *Peg3* genome browser tracks were downloaded from ENCODE. All other RNA-seq data were downloaded and processed as described in “Transcriptome assembly and annotation”. Bigwig files were generated using deeptools ^103^ (v3.5.4; bamCoverage – normalizeUsing CPM –binSize 1 –minMappingQuality 255; bigwigAverage -bs 50). Expression tracks shown counts per million of uniquely mapped reads in 200bp bins. FANTOM5 total counts were downloaded from the UCSC table browser.

To estimate the abundance of 3’-tRFs in HeLa cells, small RNA-seq data were downloaded from the sequence read archive using sra-tools (v3.0.0). Adapters were removed and trimmed reads between 14 and 50 nucleotides were retained using cutadapt ^95^ (v4.6; cutadapt -m 14 -M 50 -a TGGAATTCTCGGGTGCCAAGG). Quality filtering was performed using FASTX-Toolkit (v0.0.14; fastq_quality_filter -Q33 -q 20 -p 95) and quality was manually inspected using fastQC (v0.12.1). For alignment to tRNAs, a Bowtie2 (v2.5.3) index was constructed from *Homo sapiens* (hg38) mature tRNA sequences downloaded from [GtRNAdb](https://gtrnadb.org/genomes/eukaryota/Hsapi38/hg38-mature-tRNAs.fa) ^92^ (release 22), with “CCA” appended to each. Reads were aligned to this index using Bowtie2 (bowtie2 -N 1 -p 4 -L 10 -R 10 –gbar 20). For the purpose of calculating total mapped reads, unmapped reads (bamtools filter -isMapped false) and reads aligning with more than 2 mismatches (bamtools filter -tag XM:”>2”) were extracted and re-aligned to a tRNA-masked genome, again retaining only reads that aligned with 2 or fewer mismatches. Reads aligning to tRNAs were categorized and counted using bedtools (v2.31.1; intersectBed) followed by a custom R script. Briefly, reads were considered “tRF3b” if they ended within 3 bases of the corresponding tRNA annotation and were 21-23nt in length. Counts for each tRF3b species were mapped back to the unique IDs used for target prediction. To calculate reads per million, counts for each 3’-tRF were divided by the total number of mapped reads. Scripts used for small RNA-seq analysis are available at a [GitHub repository](https://github.com/mwpeacey/small_RNA_seq).

For analysis of *Peg3* expression in *Ago1/2*^-/-^, *Drosha*^-/-^ and *Dgcr8*^-/-^ embryonic stem cells, raw counts were downloaded from GEO using accession codes GSE122627, GSE110942, GSE78971 and GSE78974. Differential expression analysis was performed with DESeq2 (v1.48.1) ^104^, with pre-filtering for *≥*1 read count per gene in at least one sample.

### Statistical analysis

All data visualization and statistical analysis was performed in R (v4.5.1) using the packages tidyverse ^105^, patchwork, glue, pheatmap, rtracklayer ^106^ and Gviz ^107^. Odds ratios were calculated by Fisher’s exact test. Relative values were compared to a baseline of 1 by one-sided, one-sample *t* -tests. Dose response was modeled by linear regression of log_2_ relative repression on log_10_ plasmid amount, weighted by standard error. Multiple comparisons corrections were made using the Benjamini-Hochberg method. Asterisks (*) indicate a *p*-value of < 0.05. Error bars show propagated standard error of the mean from technical replicates, unless otherwise indicated.

## Acknowledgments

We thank Jenna Wilken and Samantha D’Asaro for technical assistance. This work was supported by the US National Institutes of Health grant R01 GM138669 (A.J.S.) and the George A. and Marjorie H. Anderson Fellowship (M.P.). The Cold Spring Harbor Laboratory (CSHL) NGS Sequencing Core Facility was supported by the US National Institutes of Health grant P30CA045508. Work on the CSHL high performance compute cluster was performed with assistance from the US National Institutes of Health grant S10OD028632-01.

## Author contributions

M.P. and A.J.S. designed the study; M.P. and J.I.S. performed the experiments; M.P. and A.J.S. analyzed the data and/or its significance; M.P. and A.J.S. wrote the manuscript; A.J.S. acquired funding.

## Supplementary figures

**Figure S1.**
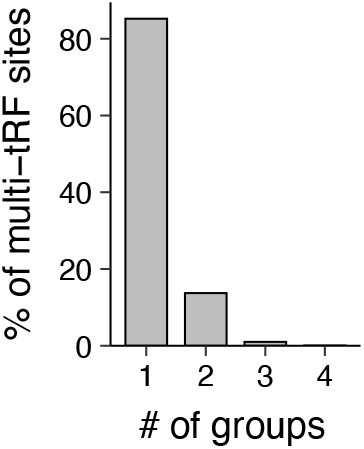
The number of tRF3b sequence groups (defined in figure 1F) per target site among sites with more than one unique tRF3b sequence aligned.

**Figure S2.**
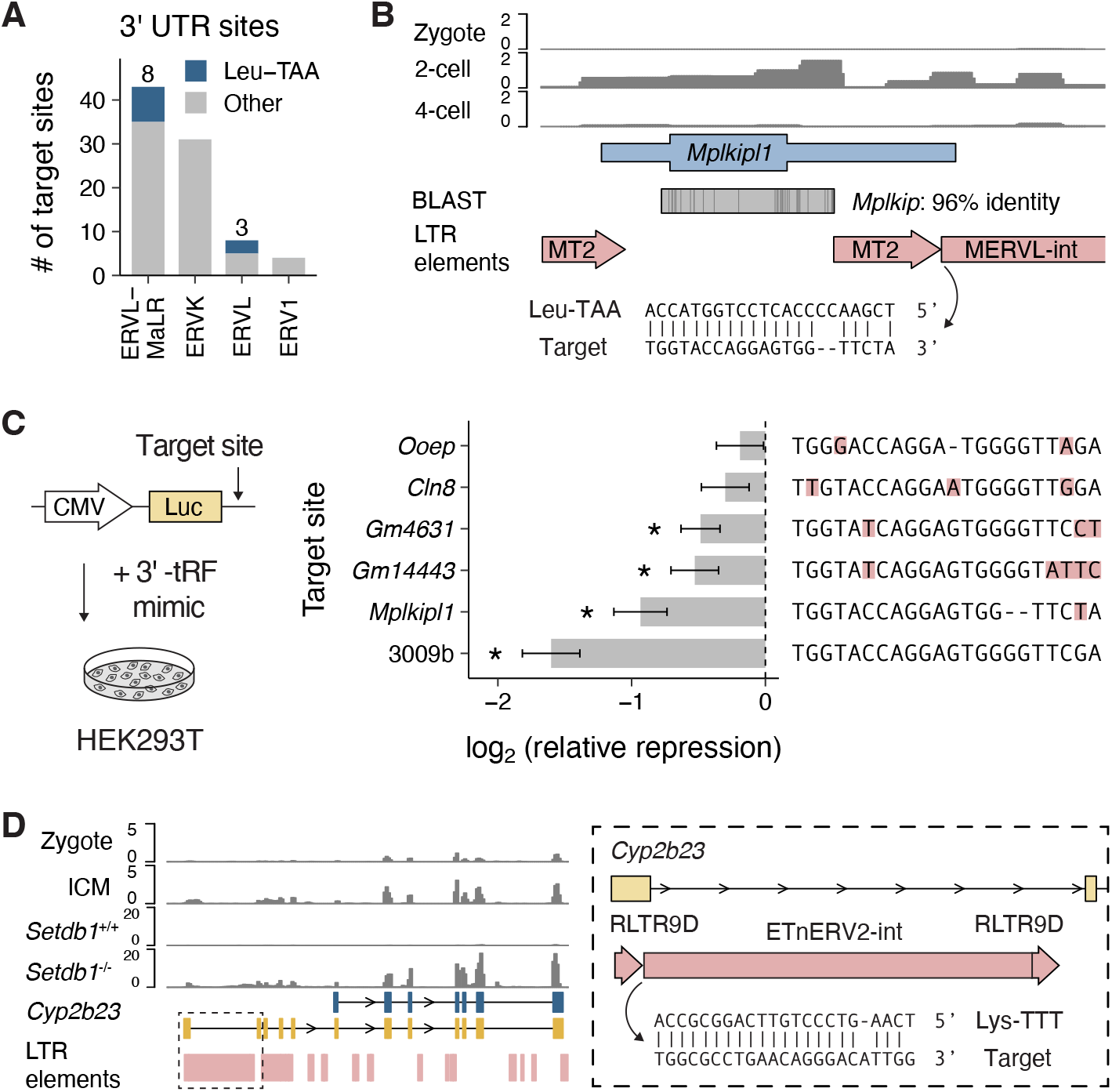
(**A**) Distribution of LTR-associated 3’-tRF target sites in the 3’ UTR of protein-coding genes across LTR-retrotransposon families. Highlighted is the number of sites for which the top scoring 3’-tRF is derived from a Leu-TAA tRNA. (**B**) Genome browser view of the *Mplkipl1* locus in the C57BL/6J reference strain. RNA-seq tracks show expression in the pre-implantation embryo (GSE66582). BLAST track shows the region of homology with *Mplkip*, with mismatches indicated in dark grey. (**C**) Repression of luciferase reporters containing 3’ UTR target sites from the indicated genes by a Leu-TAA tRF3b mimic. Relative repression was calculated by normalizing first to a no target site reporter, and then to a non-targeting tRF3b. The 3009b reporter serves as a positive control with a perfectly complementary site. For each target site, red highlighting indicates mismatches to the tRF3b sequence. Error bars show propagated standard error from technical replicates. Asterisks (*) indicate *p*-values < 0.05 from one-sided, one-sample *t* -tests. (**D**) Genome browser view of the *Cyp2b23* locus showing a StringTie assembled transcript (yellow) initiated in an LTR with a predicted 3’-tRF target site in the 5’ UTR, alongside the canonical GENCODE-annotated transcript (blue). RNA-seq tracks show expression in the pre-implantation embryo (GSE66582), and in wild-type or *Setdb1* knockout (GSE29413) mESCs.

**Figure S3.**
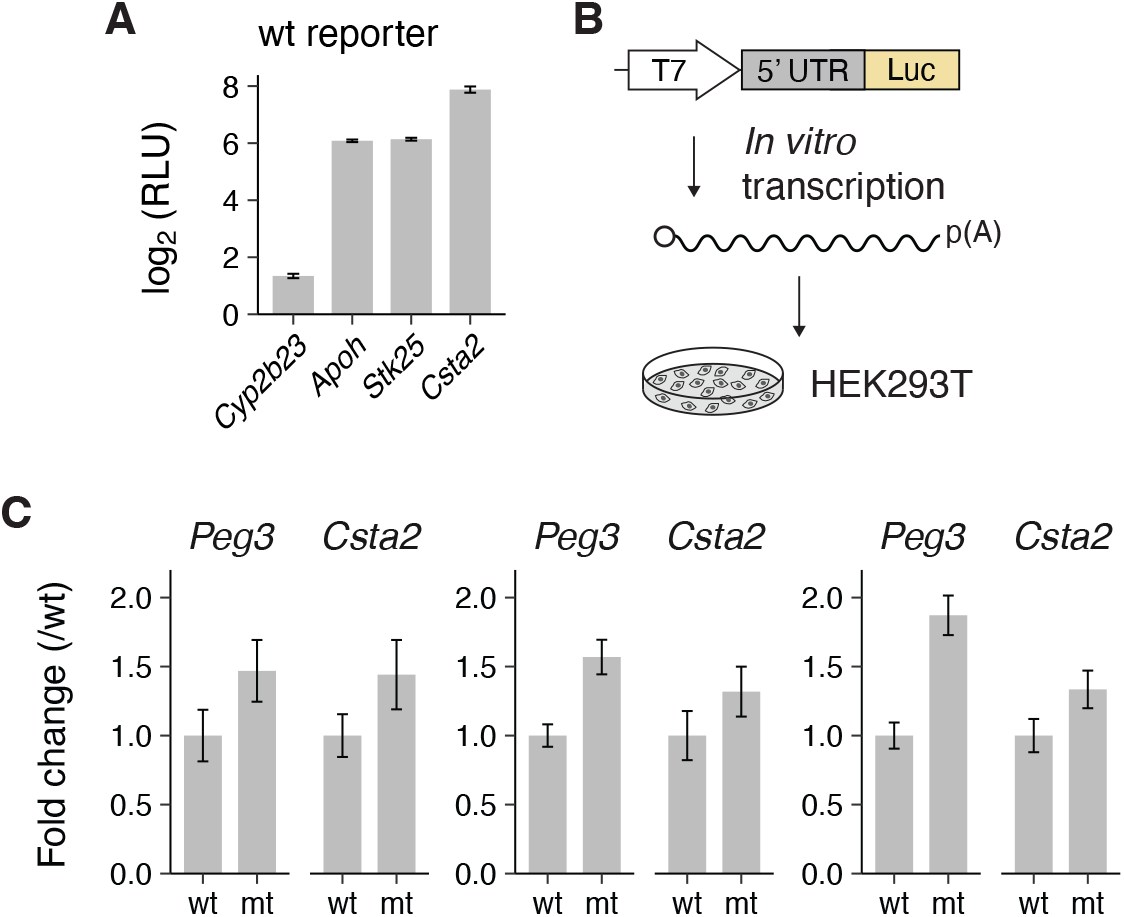
(**A**) Relative light unit (RLU) output of reporters containing the 5’ UTRs of the indicated genes with wild-type (wt) target sites. Error bars show propagated standard error from technical replicates. (**B**) Schematic of *in vitro* transcribed luciferase reporter mRNA, which is polyadenylated (pA) and includes an m7G cap (white circle). (**C**) The three biological replicates summarized in figure 3E. Error bars show propagated standard error from technical replicates.

**Figure S4.**
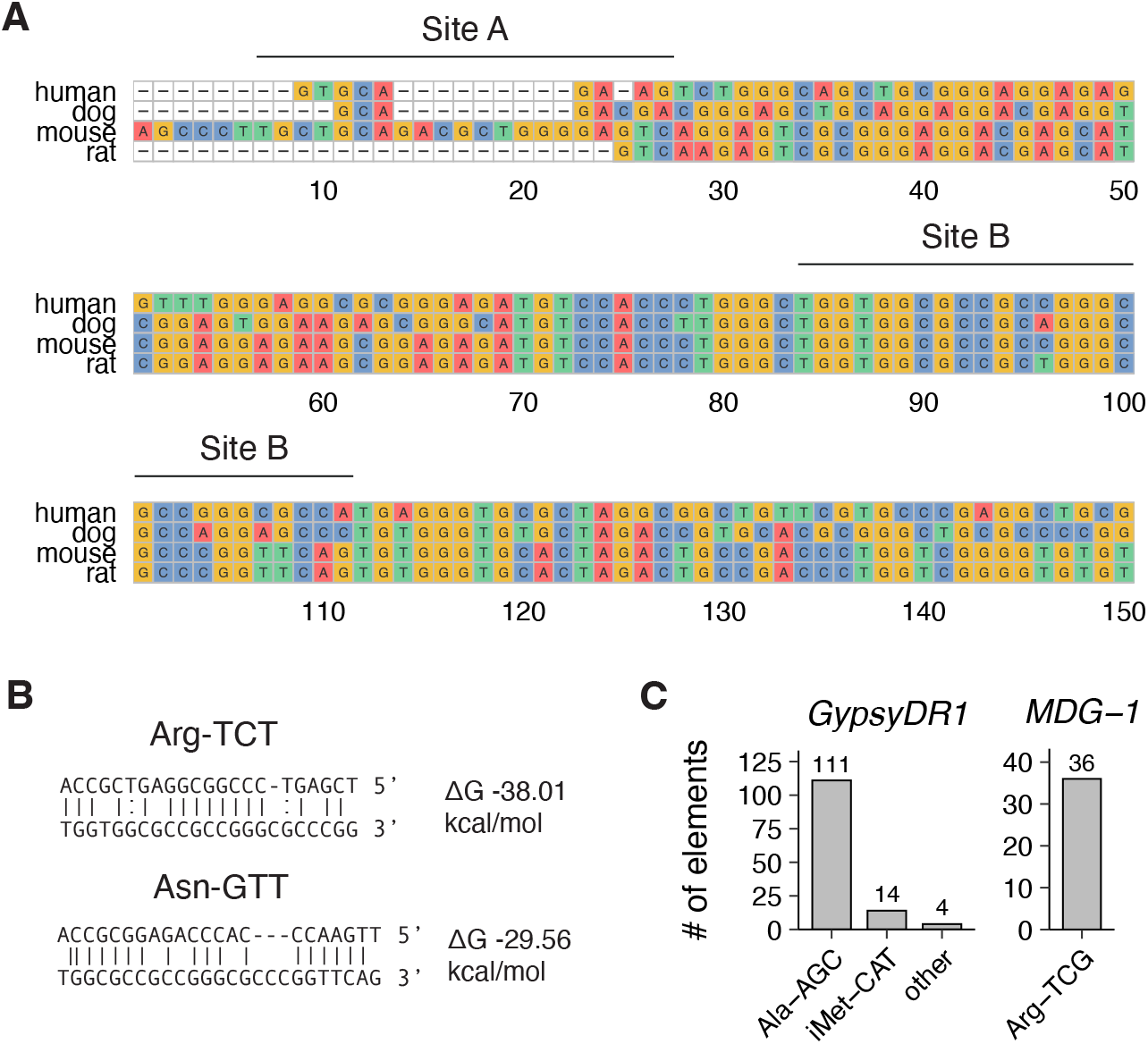
(**A**) Multiple sequence alignment of exon 1 of canonical *Peg3* transcripts across species. Predicted target sites are labeled as in figures 4A and 4B. (**B**) Alignment of top scoring tRF3b sequences to target site B in the *Peg3* 5’ UTR, with associated Gibbs free energies of interaction (ΔG). (**C**) Priming tRNAs assignable to *GypsyDR1* elements in *Danio rerio* (left; danRer11) and *MDG-1* elements in *Drosophila melanogaster* (right; dm6).

